# Fast genome-based species delimitation: Enterobacterales and beyond

**DOI:** 10.1101/2023.04.05.535762

**Authors:** Julie E. Hernández-Salmerón, Tanya Irani, Gabriel Moreno-Hagelsieb

## Abstract

Average Nucleotide Identity (ANI) is becoming a standard measure for bacterial species delimitation. However, its calculation can take orders of magnitude longer than fast similarity estimates based on sampling of short nucleotides, compiled into so-called sketches. These estimates are widely used and correlate well with ANI. However, they might not be as accurate. Thus, we compared two sketching programs, mash and dashing, against ANI, in delimiting species among publicly available Esterobacterales genomes. Receiver Operating Characteristic (ROC) curve analysis found all three programs to be highly accurate, with Area Under the Curve (AUC) values of 0.99, indicating almost perfect species discrimination. Subsampling to reduce over-represented species, reduced these AUC values to 0.92. Focused tests with ten genera represented by more than three species, also showed almost identical results for all methods. *Shigella* showed the lowest AUC values (0.68), followed by *Citrobacter* (0.80). All other genera, *Dickeya, Enterobacter, Escherichia, Klebsiella, Pectobacterium, Proteus, Providencia* and *Yersinia*, produced AUC values above 0.90. The species delimitation thresholds varied, with species distance ranges in a few genera overlapping the genus ranges of other genera. Mash was able to separate the *E. coli* + *Shigella* complex into 25 apparent phylogroups. Testing mash for species separation in genera outside Enterobacterales showed AUCs above 0.95, again with different thresholds for species delimitation within each genus. Overall, our results suggest that fast estimates of genome similarity are as good as ANI for species delimitation. Therefore, these fast estimates might suffice for determining the role of genomic similarity in bacterial taxonomy.

## 1. Introduction

Average Nucleotide Identity (ANI), seems to be becoming a standard in genome sequence-based species delimitation (6, 11, 14, 17). The method involves the comparison of genome segments, around 1000 base-pairs long, against another genome, and, as the name implies, the calculation of the average identity of matching segments. While much faster than experimentally-based approaches, such as DNA-DNA hybridization, the method can take a very long time to compare thousands of genomes, even when using an optimized program for the calculation, such as fastANI (8).

Methods based on sampling k-mers, normally between 20 and 40 bp long, can produce genome similarity/distances orders of magnitude faster than ANI (3, 7, 13). The most commonly used of these methods is the MinHash approach, programmed into the mash software (13). This program gathers k-mer samples, codified as hashes, into so-called sketches, which can be efficiently compared. The overall genome similarity is then estimated from the proportion of identical kmers found between the compared sketches. MinHash can estimate genome similarities in a fraction of the time required for ANI (7, 13), and have been reported to highly correlate with ANI measurements (13). Another program, dashing (3), offers a few different approaches to codifying sketches, and can produce results very similar to those by mash, among other useful genome similarity estimates (3).

While the articles on mash and dashing show that their results correlate very well with those produced by ANI (3, 13), more where-the-rubber-meets-the-road analyses seem to be required to properly evaluate how well they would substitute overall genome similarity comparisons. We recently published a comparison of these programs against ANI in delimiting species in the *Klebsiella* genus (7). Since the classification of *Klebsiella* has involved ANI along other features (15, 19, 20), it is not surprising that ANI-based clusters had an excellent match with species annotations in this genus. The good news was that the sketch-based methods produced results of the exact same quality (7).

Here we present an expansion of those analyses towards all of the complete genomes available at NCBI’s RefSeq genome database (11), classified into the Enterobacterales taxonomic order, to try and ensure that the sketch-based estimates can work as well as ANI in delimiting species beyond those of *Klebsiella*. We therefore tested fastANI, mash and dashing for delimiting species at the overall order level, as well as within genera represented by more than two species.

## 2. Methods

### A. Genome sequences

To compile a genome sequences dataset, we downloaded all available genomes with a “Complete” status at NCBI’s RefSeq genome database (11), by the middle of January of 2023. From these, we selected all the genomes classified into the order Enterobacterales, as long as they were classified to the species level, for a total of 7,132 genome sequences (Supplementary Table S1). Type strains were identified from the annotations found in the corresponding NCBI’s “GBFF” files.

### B. Genome comparisons

To calculate Average Nucleotide Identity (ANI), we selected the fastANI program (8), produced by the group that, as far as we know, has most promoted the use of ANI in genomewide classification studies (8–10, 22). Briefly, the original ANI calculations (5), started by breaking a query genome into 1020 base-pair long segments, which were then compared against a target genome using NCBI’s blastn (2). As the name suggests, ANI was then calculated by taking the average identity of segments matched by blastn. Instead of blastn, fastANI uses a MinHash approach to quickly estimate the similarity between the genome-derived DNA segments and the target genome (8). To run fastANI comparisons, we chose a fragment length of 1020 bp, instead of the default of 3000 bp (--fragLen 1020). Every other option was left at its default value. The ANI produced by fastANI is reported as a percent identity. Therefore, to transform ANI into distances, we subtracted the obtained ANI from 100 and divided the result by 100.

To calculate distances using the MinHash approach, we used two programs: mash (13) and dashing (3). For mash, version 2.3, we produced sketches of 5000 k-mers (-s 5000 option), rather than the default of 1000, because our preliminary tests showed that the default failed to produce distances for many genomes. We did not use more than 5000 sketches, because our preliminary tests also found that the results were almost identical to those produced using 10,000 k-mers (data not shown). Every other option was left to its default value. The distances produce by mash are reported as fractions, rather than percents.

We used dashing version 1.0.2. We changed the k-mer length to 21 bp, instead of its default of 31, for two reasons: (1) because 21 is the k-mer size used by mash, and (2) with 31 bp, the program failed to produce estimated distances for several genome comparisons. We used a sketch size of 2^1^4 (-S 14), because this sketch size produced the best estimates of genomic Jaccard similarity in the original dashing publication (3). We only tested mash distances (-M option) because our prior report showed that results from other measures were identical (7). Every other option was left unchanged from its default value.

### C. Accuracy and plotting

Plots and other calculations were produced using R (16). To test the accuracy of all programs, we performed Receiver Operating Characteristic (ROC) curve analyses as implemented in the R package cutpointr (21). We also used the cutpointr package to find optimal cutoffs. Trees were displayed using the R packages ggtree (24) and ggtreeExtra (23).

## 3. Results and Discussion

Though the focus of this report is on the quality of results, it is noteworthy that calculating ANI between all genomes, with the fastANI program, required the separation of all genomes into several datasets to divide the labour across three computers. The calculation took around two months. Calculating distances using mash or dashing took no more than a couple of hours, including the time to produce “presketch” files, which can be reused to produce results in a more timely manner. These fast programs were run in an iMac computer with 32G of RAM and an Intel processor. The programs ran using four of the available 8 cpu threads.

### A. All methods discriminated Enterobacterales species with high accuracy

To test species/genus separation using all three tools, fastANI, mash and dashing, we produced tables with all *vs*. all pairwise distances, as calculated with all three methods (Supplementary Table S2). Pairs belonging to either the same species, or the same genus / different species, were used to test for species discrimination. A test with all same species and same genus pairs showed AUC values above 0.99 for all three methods (Fig. 1).

**Fig. 1.**
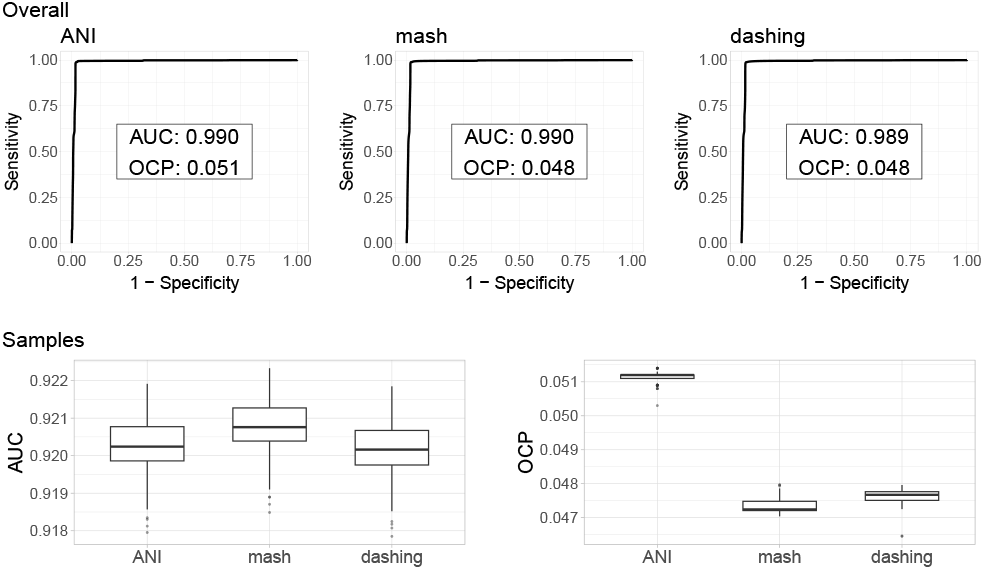
Species separation across Enterobacterales. All three methods produced exactly the same overall quality of species separation with Area Under the Curve (AUC) values close to 0.99. Optimal cutoff (OCP) points for species separation suggested by the analysis are somewhat different though. After subsampling genomes from over-represented species, to avoid bias, the AUC lowered to a median slightly above 0.92 with all programs, with OCP similar to those obtained with all the genomes.

Prior results suggested that minHash, represented by mash, was not as accurate as ANI (8). However, that comparison was about whether the results correlated well with ANI as traditionally calculated, rather than about actual accuracy in the task of delimiting species (8).

The species representation of the Enterobacterales genome dataset is heavily biased. The dataset contains a number of *Escherichia coli* representatives orders of magnitude larger than those of most other species. Therefore, the AUC values might be biased towards the distinction of *E. coli* (of all 4,381,875 same species pairs, 2,671,516, or 61%, are *E. coli* pairs). The abundance of *E. coli, Klebsiella pneumoniae* (944,625 pairs, 22%) and *Salmonella enterica* (697,971 pairs, 16%) covered 99% of all same species pairs, even though the dataset contained 256 species, with 139 of them represented by more than one genome.

To avoid the effects of these biases, we obtained 100 subsamples each containing 100 genomes randomly sampled from all species represented by more than 100 genomes. This leveled the number of the top five most represented species. All other species kept their total genomes, with several of them having representative numbers within the same order of magnitude of the subsamples. Again, all three methods produced results of the same quality, though the median AUC values were slightly above 0.92 (Fig. 1), lower than the AUC values obtained in the naïve test above, but still indicating very good accuracy in species separation.

### B. Different genera showed different within-species distances with all methods

The tests above combine all same species and same genus pairs, regardless of potential for confusion. Since the most difficult distinction should be between genomes from organisms of the same species from those in the same genus, we ran further tests choosing genera from the Enterobacterales, represented by at least three species, making sure that all three species within the selected genera were represented by ten genomes or more. This process resulted in ten genera: *Citrobacter, Dickeya, Enterobacter, Escherichia, Klebsiella, Pectobacterium, Proteus, Providencia, Shigella*, and *Yersinia*.

Of the ten genera above, two had severe biases in abundance of genomes representing a single species: *Escherichia* and *Klebsiella*. These were subject to subsampling to the number of genomes in the second most abundantly represented species.

The resulting AUC values for species discrimination between these genera varied (Table 1). The lowest values obtained were for *Shigella*, with AUC values of approx. 0.68. That value was followed by 0.80 for *Citrobacter*. Most other AUC values were above 0.90 (Table 1). Again, the results were pretty much identical with all three measures.

**Table 1.** *The values for *Escherichia* and *Klebsiella* are the median AUC calculated from 20 sub-samples.

| Area under the curve (AUC) values for species/genus separation across ten genera. |  |  |  |  |
| --- | --- | --- | --- | --- |
| Genus | No. species | ANI | mash | dashing |
| *Citrobacter* | 14 | 0.805 | 0.802 | 0.804 |
| *Dickeya* | 9 | 0.999 | 0.999 | 0.999 |
| *Enterobacter* | 14 | 0.921 | 0.921 | 0.922 |
| *Escherichia* * | 4 | 1.000 | 1.000 | 1.000 |
| *Klebsiella* * | 11 | 0.987 | 0.988 | 0.987 |
| *Pectobacterium* | 16 | 0.964 | 0.965 | 0.963 |
| *Proteus* | 6 | 0.995 | 0.995 | 0.995 |
| *Providencia* | 7 | 0.971 | 0.972 | 0.954 |
| *Shigella* | 4 | 0.690 | 0.673 | 0.677 |
| *Yersinia* | 17 | 0.990 | 0.990 | 0.989 |

*Klebsiella* is the genus with the best species representation of all ten, with seven species represented by more than ten genomes, six of them by more than 30. *K. pneumoniae* exceeded the representation of the next more represented species, *K. quasipneumoniae* by an order of magnitude. We therefore produced 50 subsamples topping 50 representative genomes per species, thus leveling the order of magnitude in representation for the top six genomes. The AUC for species discrimination in this genus remained close to 0.99 in all subsamples with all programs tested (Table 1). As explained above, ANI has been part of the taxonomic assignments in the genus (15, 19, 20), which explains the high accuracy obtained with this measure. It is still important, however, to note that mash and dashing produced results of the same quality. We have presented a more focused work about species delimitation in this genus before (7).

The *Escherichia* genus also had a top species, *E. coli*, represented by orders of magnitude more genomes than the next one, *E. fergusonii*. We therefore produced 50 subsamples of the genus, topping the number of representative genomes per species to 50, thus leveling the order of magnitude of three of the four species in this genus. The separation between these species was always clean (AUC: 1.00).

Distance ranges varied among the species in the ten genera (Fig. 2). *Yersinia* showed the lowest values, while *Providencia* showed species values overlapping the within-Genus distributions of most other genera. This variation agrees with a previous study focused on the ANI distributions of several bacterial species (12), but contradicting the notion of a universal genetic species delimitation (8, 18). This result also seems to conflict with the high accuracy for species delimitation across all Enterobacterales that we presented above. However, the apparent conflict is resolved by the fact that the species in seven of these genera have distance ranges below, roughly, 0.05, combined with within-genus distance ranges above the same mark.

**Fig. 2.**
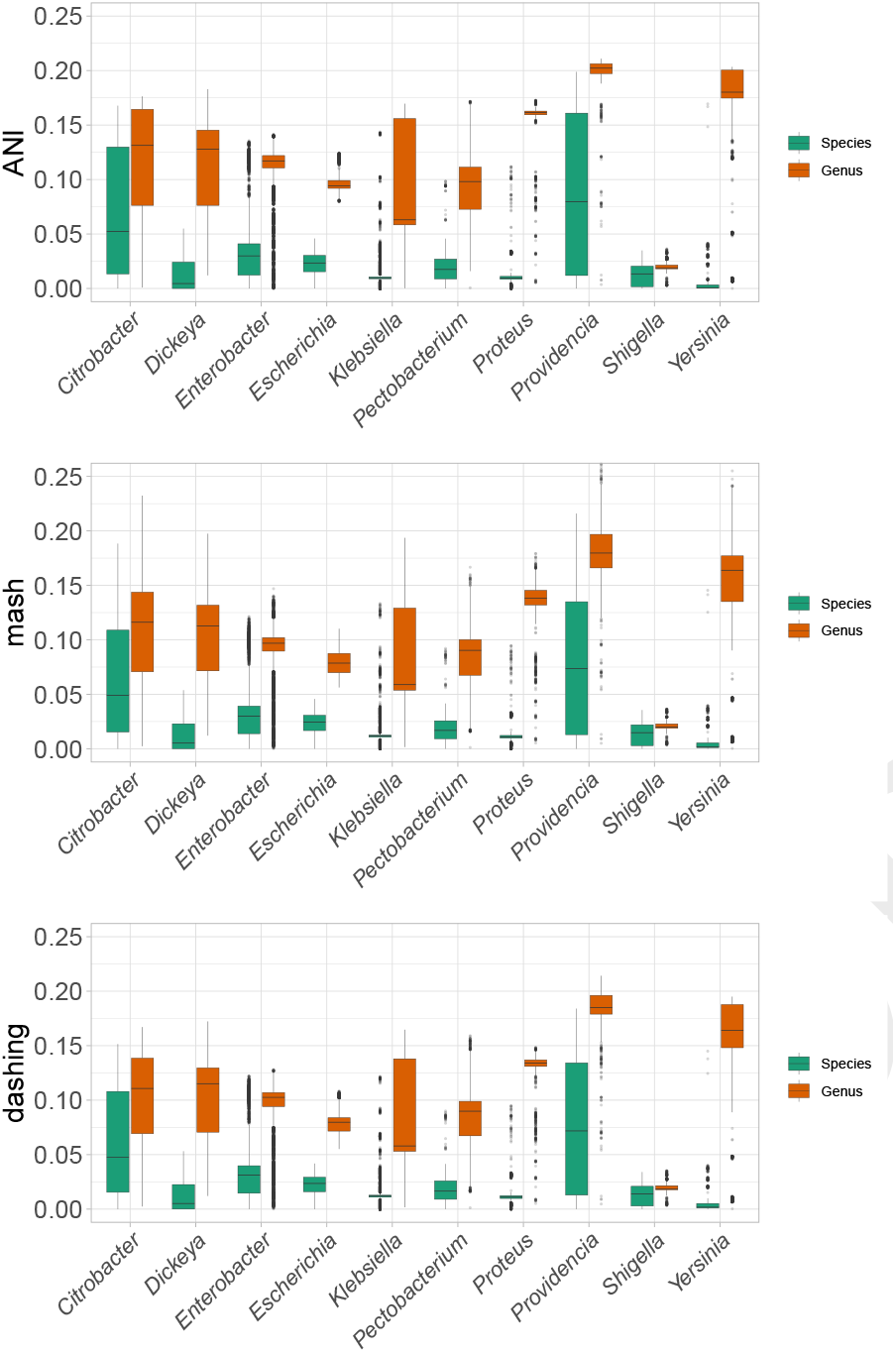
Species distances. The range of species distances varied from one genus to the next, suggesting that there is no universal species delimitation. Though results are very similar for all measures, dashing seemed to produce more outliers.

### C. The *E. coli* + *Shigella* complex separated into 25 phylogroups

The separation of species in the *Escherichia* genus was clean for all samples tested (AUC: 1), while that for *Shigella* was the worst (AUC: 0.68, see above). However, the clustering of *Shigella* within the *E. coli* species is a classic taxonomy *vs*. tradition issue (discussed in 4). Accordingly, cutting hierarchical clusters at an ANI distance of 0.051, or a mash distance of 0.048 (the cutoffs suggested for the whole dataset when optimising for the F1 score using the cutpointr package; Fig. 1), produces a cluster containing all *E. coli* genomes combined with all the genomes representing the four *Shigella* species in the dataset (Fig. 3). No other species belonging to different genera showed such within species distances in our dataset. We examined this group more closely.

**Fig. 3.**
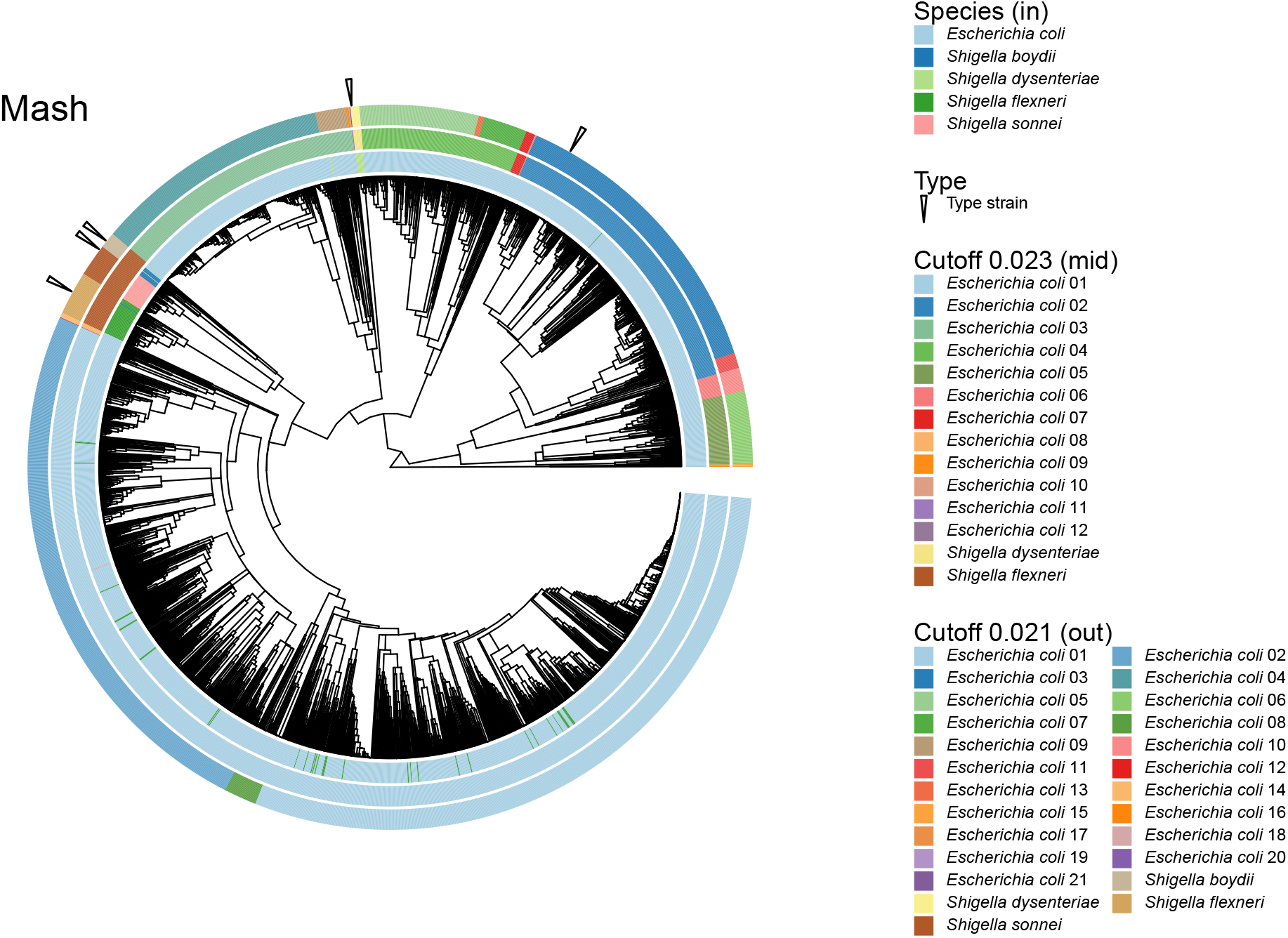
The *Escherichia coli* + *Shigella* complex. The hierarchy shows that several *Shigella* genomes clustered together forming coherent groups, though many are interspersed into groups dominated by genomes labeled as *E. coli* (inner circle, or circle closest to the dendogram). A mash distance cutoff of 0.023 produced 14 groups, two of them mainly composed of *Shigella* genomes (middle circle). A lower cutoff, 0.021, separated *Shigella* into all four represented species, but it also divided the whole group into a total of 25 clusters (external circle), or apparent phylogroups.

**Fig. 4.**
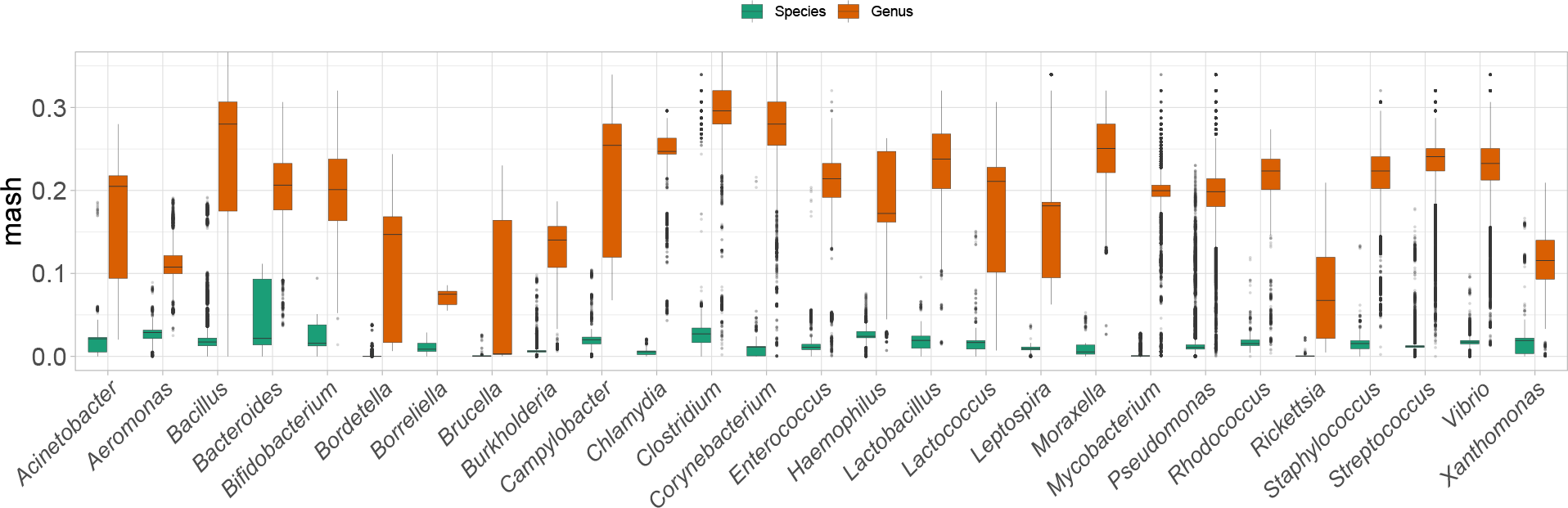
Species distances outside Enterobacterales. The range of species distances also varied from one genus to the next, suggesting that there is no universal species delimitation across bacteria.

Examination of the hierarchical cluster produced from either distance measure showed some clean groups composed of *Shigella* species (Fig. 3, inner circle). However, other *Shigella* strains were interspersed with clear *E. coli* groups. Similarly, a few genomes annotated as *E. coli* were inserted into *Shigella* groups. The latter two cases apparently reveal mislabeled genomes. Therefore, it is possible that the anomalously clustered genomes need to be relabeled according to the main names within their apparent phylogroups. To test for this possibility, we looked for a mash distance cutoff that could cleanly separate the observed clusters that coincide with *Shigella* species. A mash distance cutoff of 0.021 worked for this purpose. In other words, this cutoff produced four *Shigella* clusters, each corresponding to a *Shigella* species, and each containing their corresponding type strain (Fig. 3, outermost circle; Supplementary Table S3). Two of these were clean single-species groups: *S. sonnei* and *S. dysenteriae*. The *S. flexneri* group contained an apparently mislabeled *S. boydii* genome, while the *S. boydii* group contained five *E. coli* genomes and two *S. dysenteriae* ones. This cutoff produced 21 *E. coli* groups, for a total of 25 apparent phylogroups (Fig. 3, outermost circle; Supplementary Table S3). Of these, 17 were clean *E. coli* clusters. Three of the four apparently contaminated *E. coli* groups were mixed with *S. flexneri*, with one of them also containing a genome labeled as *S. sonnei*. The remaining *E. coli* group contained two genomes labeled as *S. dysenteriae*.

A prior analysis, founded on mash distances and supported by further analyses, suggested that the *E. coli* + *Shigella* complex, could be divided into 14 phylogroups (1), fewer than our suggested 25 phylogroups above. Therefore, we found a mash distance cutoff that would cut the hierarchical cluster into 14 groups (0.023). Only two of these were *Shigella* groups, in apparent agreement with the two *Shigella* phylogroups suggested by (1), though our two *Shigella* groups were separated in the hierarchy by clean *E. coli* groups (Fig. 3, mid circle; Supplementary Table S3), rather than next to each other as in the prior study (1). One of the *Shigella* groups was mainly composed of genomes named after all four species: *S. flexneri, S. sonnei, S. boydii* and *S. dysenteriae*, in order of abundance (Fig. 3, mid and outermost circles; Supplementary Table S3). The second *Shigella* cluster was composed exclusively of ten *S. dysenteriae* genomes. Three of the remaining twelve *E. coli* groups contained genomes named after *Shigella* species (Fig. 3, mid circle; Supplementary Table S3).

The results presented in this section suggest that, though most genomes showed phylogenomic coherence matching their species labels, some genomes in the group were mislabelled. Though our 25 apparent phylogroups seemed coherent, deciding whether the *Shigella* clusters should correspond to two or more phylogroups might require further analyses that are beyond the scope of this research.

### D. Species delimitation with mash for genera belonging to other taxonomic orders showed AUCs above 0.95

Convinced that mash and dashing produce results of the same quality as ANI, we proceeded to examine species delimitation in genera outside Enterobacterales using mash. We selected these genera by the same criteria as above. Namely, these genera had at least three species represented by at least ten genomes in the database. The procedure yielded 27 genera (Table 2).

**Table 2.** *The values for *Acinetobacter, Bordetella, Enterococcus, Pseudomonas*, and *Staphylococcus* are median AUCs calculated from 50 subsamples.

| Genus | Order | No. species | AUC |
| --- | --- | --- | --- |
| *Aeromonas* | Aeromonadales | 15 | 0.999 |
| *Bacillus* | Bacillales | 49 | 0.975 |
| *Staphylococcus* * | Bacillales | 43 | 1.000 |
| *Bacteroides* | Bacteroidales | 21 | 0.988 |
| *Bifidobacterium* | Bifidobacteriales | 22 | 1.000 |
| *Bordetella* * | Burkholderiales | 11 | 0.996 |
| *Burkholderia* | Burkholderiales | 32 | 0.993 |
| *Campylobacter* | Campylobacteriales | 35 | 1.000 |
| *Chlamydia* | Chlamydiales | 15 | 1.000 |
| *Corynebacterium* | Corynebacteriales | 91 | 1.000 |
| *Mycobacterium* | Corynebacteriales | 59 | 0.999 |
| *Rhodococcus* | Corynebacteriales | 16 | 0.985 |
| *Clostridium* | Eubacteriales | 40 | 0.953 |
| *Brucella* | Hyphomicrobiales | 15 | 0.982 |
| *Enterococcus* * | Lactobacillales | 16 | 0.998 |
| *Lactobacillus* | Lactobacillales | 21 | 0.999 |
| *Lactococcus* | Lactobacillales | 12 | 0.996 |
| *Streptococcus* | Lactobacillales | 62 | 1.000 |
| *Leptospira* | Leptospirales | 11 | 1.000 |
| *Acinetobacter* * | Moraxellales | 38 | 0.998 |
| *Moraxella* | Moraxellales | 8 | 1.000 |
| *Haemophilus* | Pasteurellales | 7 | 0.978 |
| *Pseudomonas* * | Pseudomonadales | 146 | 0.965 |
| *Rickettsia* | Rickettsiales | 21 | 0.989 |
| *Borrelia* | Spirochaetales | 5 | 1.000 |
| *Vibrio* | Vibrionales | 86 | 1.000 |
| *Xanthomonas* | Xanthomonadales | 21 | 0.994 |

The AUC values for species delimitation were above 0.95 for all of these genera, with several of them showing perfect discrimination (AUC: 1.00) (Table 2). Of the 27 genera examined, five had a top species represented at least an order of magnitude above the next most represented species: *Acinetobacter, Bordetella, Enterococcus, Pseudomonas*, and *Staphylococcus*. In these cases, we produced 50 subsamples limiting the number of representative genomes per species to 50. As with the Enterobacterales genera presented above, the AUC values in Table 2 are medians as obtained from the 50 samples.

The AUC analyses also suggested different optimal species/genus cutpoints among the examined genera. Therefore, we checked the species and genus mash distance ranges (Fig.4; Supplementary Table S4). As was the case for the Enterobacterales genera, the species mash distance ranges varied from genus to genus. As above, the species distance ranges of some genera overlapped the genus ranges of other genera, suggesting that a single mash distance threshold might not work well for some cases.

## 4. Conclusion

Overall, our results suggest that using fast, sketch-based estimates of genome similarity, can be as accurate for bacterial taxonomy as whole-genome Average Nucleotide Identity. Therefore, these estimates should suffice for further determination of the role that genome similarity should play in bacterial taxonomy.

## Acknowledgements

This work was supported by the Natural Sciences and Engineering Research Council of Canada (NSERC) discovery grant: RGPIN:2018-06180.

